# Supervised learning on synthetic data for reverse engineering gene regulatory networks from experimental time-series

**DOI:** 10.1101/356477

**Authors:** Stefan Ganscha, Vincent Fortuin, Max Horn, Eirini Arvaniti, Manfred Claassen

## Abstract

The reconstruction of gene regulatory networks from time resolved gene expression measurements is a key challenge in systems biology with applications in health and disease. While the most popular network inference methods are based on unsupervised learning approaches, supervised learning methods have proven their potential for superior reconstruction performance. However, obtaining the appropriate volume of informative training data constitutes a key limitation for the success of such methods.

Here, we introduce a supervised learning approach to detect gene-gene regulation based on exclusively synthetic training data, termed *surrogate learning*, and show its performance for synthetic and experimental time-series. We systematically investigate different simulation configurations of *biologically representative* time-series of transcripts and augmentation of the data with a measurement model. We compare the resulting synthetic datasets to experimental data, and evaluate classifiers trained on them for detection of gene-gene regulation from experimental time-series. For classifiers, we consider hybrid convolutional recurrent neural networks, random forests and logistic regression, and evaluate the reconstruction performance of different simulation settings, data pre-processing and classifiers.

When training and test time-courses are generated from the same distribution, we find that the largest tested neural network architecture achieves the best performance of 0.448 ± 0.047 (mean ± std) in maximally achievable F1 score over all datasets outperforming random forests by 32.4 % ± 14 % (mean ± std). Reconstruction performance is sensitive to discrepancies between synthetic training and test data, highlighting the importance of matching training and test data domains. For an experimental gene expression dataset from *E.coli*, we find that training data generated with measurement model, multi-gene perturbations, but without data standardization is best suited for training classifiers for network reconstruction from the experimental test data. We further demonstrate superiority to multiple unsupervised, state-of-the-art methods for networks comprising 20 genes of the experimental data from *E.coli* (average AUPR best supervised = 0.22 vs best unsupervised = 0.07).

We expect the proposed surrogate learning approach to be broadly applicable. It alleviates the requirement for large, difficult to attain volumes of experimental training data and instead relies on easily accessible synthetic data. Successful application for new experimental conditions and other data types is only limited by the automatable and scalable process of designing simulations which generate suitable synthetic data.

## 1 Introduction

Gene regulatory networks constitute a central cellular information processing system and play a key role in defining health and disease states [29]. The introduction of genome-wide transcriptomic measurements opened the opportunity to reconstruct gene regulatory networks at a genome-wide scale [14]. Reconstruction
 of gene regulatory networks has proven to be a difficult task, has been addressed for different experimental designs, by a variety of computational analysis approaches, and is a target of ongoing research [38].

The most popular techniques for gene regulatory network reconstruction take an unsupervised approach. In general, these methods explicitly or implicitly assume models for gene regulation, such as stochastic processes or dynamic system models. They then derive metrics for the assessment of gene regulation from the observed gene expression measurements. They predict edges according to partial correlation and mutual information between genes or, for regression-based approaches, predict the expression levels of individual genes from measurements of other genes, and interpret the sparse coefficients as regulation [44]. Concretely, *GENIE3* (random forest regression) [31], *Context likelihood of relatedness* (CLR), a statistical approach, [17] and the *Inferelator*, based on mechanistic, ordinary differential equations [7], are well-established, unsupervised methods, all of which achieved good performance in the DREAM gene regulatory network inference challenges [50, 43, 44]. Recent approaches, tailored explicitly for time-series data, include *dynGENIE3* [30], an extension of the aforementioned *GENIE3*, and a LASSO based approach, integrating multiple datasets of time-series [47].

Gene regulatory network inference has also been cast as a supervised learning problem [64]. Such approaches learn patterns for assessment of gene regulation from data with known gene regulation relationships, in contrast to the unsupervised approaches above, which operate on metrics from models of gene regulation. Supervised gene regulatory network inference requires sufficient labeled training data, e.g. individual measurements for pairs of genes and their regulatory relationships. Supervised learning methods, such as random forests or support vector machines, can be trained to predict regulatory relationships from the time-series data. *SIRENE* [45] performs local binary classification by training support vector machines on known interactions of single transcription factors in experimental data, and predicts novel regulated genes. *CompareSVM* [22] evaluates the performance of different SVM kernels in order to predict gene regulation in synthetic data and in [60] Kernel-PCA is used to infer novel regulatory edges from time-series data. Semi-supervised learning with SVMs and random forests is performed in [48] on synthetic and real data. While neural networks have not been proposed for classification of gene expression time courses, these have been utilized to analyze sequential data in multiple other application domains [49, 27, 58, 55], in particular using Recurrent Neural Networks (RNNs), such as Long Short-Term Memory (LSTM) [28]. Applications to time-series analysis [37, 21] include Fully Convolutional Networks (FCN) and Residual networks [68], multi-channel deep convolutional neural networks [69] and Attention LSTMs and FCNs [34]. Additionally, several approaches utilized recurrent neural networks in order to describe the temporal evolution of biochemical species mechanistically and infer regulatory edges from the learned weights of the neural networks’ nodes. (See [65, 40, 51] and references therein.)

Supervised methods have achieved good performance when appropriate volumes of training data are available. If this condition is not met, due to limited availability and labelling of experimental training data, *representative* synthetic data can be utilized for training, for example in computer vision [25, 32, 63]. Here, we propose the use of synthetic data of gene expression dynamics for classifier training, circumventing the difficulties associated with the low availability of appropriate data. The resulting classifier is then utilized to reconstruct gene regulatory networks from the scarce experimental data. By simulation, the amount of synthetic training data can be effectively scaled up to arbitrary levels, but necessitates in exchange the generation of data, which is representative of the observed biological process and measurement. General mechanisms and dynamic modeling of intra-cellular, biochemical processes have been extensively studied [66], allowing for the simulation of biologically representative data [41, 23, 35, 11]. In addition, the technical variability of measurement processes has been explored empirically and formalized in a way applicable for forward simulation, for example for microarrays [61, 36] or scRNA-seq data [70, 2].

These considerations motivate a supervised learning approach for gene regulatory network reconstruction. We benchmark and assess the importance of the main conceptual components of this approach: (1) the simulation of representative data (2) the adaption of the simulations for our specific experimental dataset and (3) supervised learning. From a transfer learning point of view, we design the distribution of the source data to be similar to that of the target such that no further adaptation of the classifier training is necessary. We term this procedure of generating synthetic data, training supervised classifiers on it and applying them to experimental data *surrogate learning* (see overview in Fig. 1).

**Figure 1.**
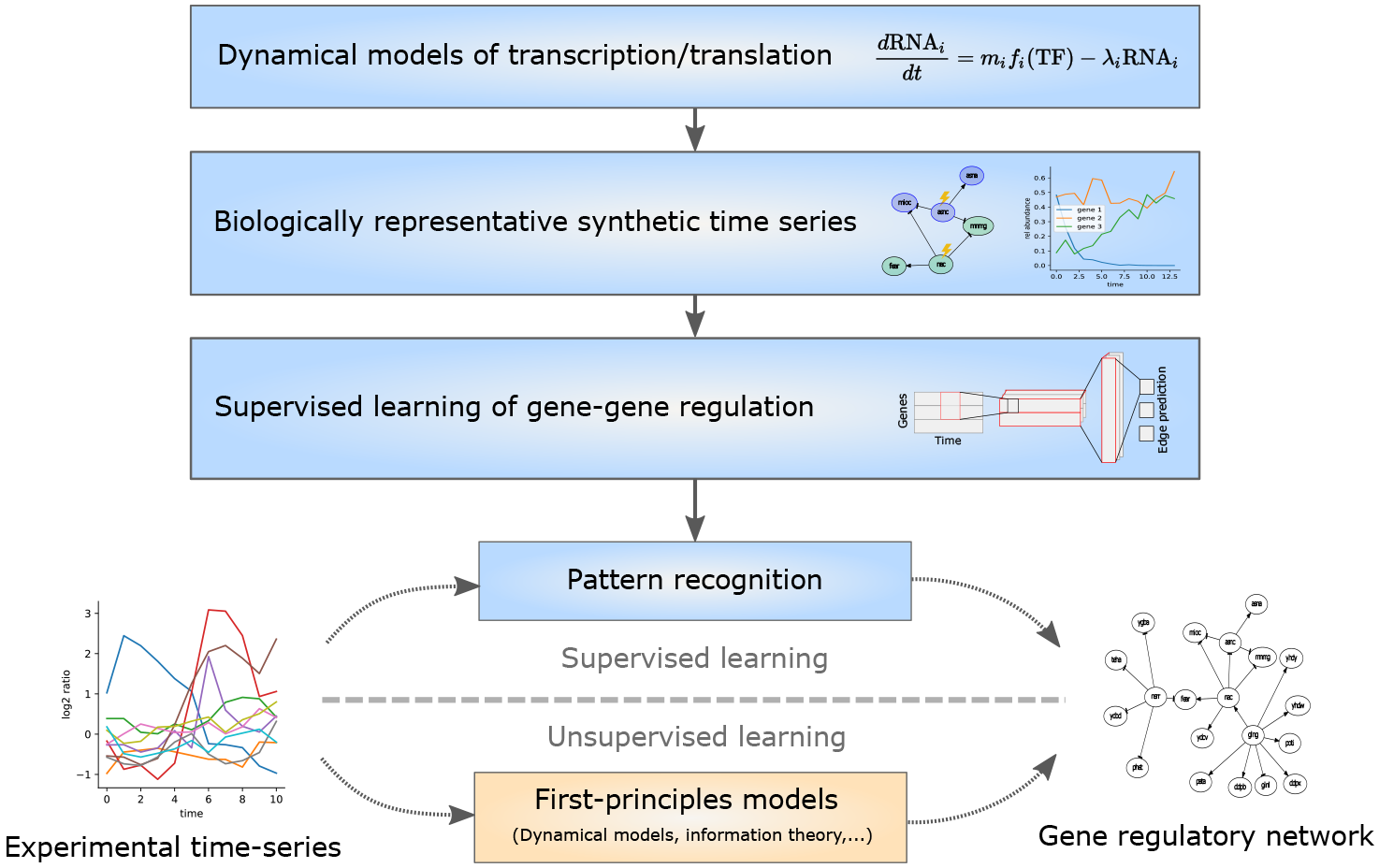
Overview of *surrogate learning* approach for gene regulatory network inference. From curated whole genome transcription factor-gene networks we extract subnetworks of size 20 and simulate them with random, but biologically representative dynamics, including different perturbation settings. We extract informative 2/3-tuples of genes from the resulting time-courses to train classifiers (neural networks, random forests) for network reconstruction. The trained classifiers are subsequently used to reconstruct gene regulatory networks from experimental time series data.

## 2 Results

We introduce a supervised learning approach for gene regulatory network inference, namely of predicting directed gene-gene interactions from time-series data, demonstrated in this study with transcriptomic bulk measurements. For this purpose, we create synthetic, but *biologically representative* transcriptomic data by simulating transcription, translation and genetic regulation for actual biological network structures and random kinetic parametrizations. Subsequently, we train classifiers on this data to reconstruct the simulated gene-gene interactions, and then utilize them to reconstruct such interactions from (possibly small-scale) experimental studies.

### 2.1 Simulation of biologically representative perturbation time series data

Data simulation aims to generate a set of biologically representative time course measurements of transcripts under perturbation, covering a wide range of biologically possible behaviours. These simulations must account for variability induced by the biological processes, as well as by measurement. For our study, we focused on microarray measurements of *E. coli* transcripts, due to the availability of time course data [5] and the large volume of prior knowledge on this species’ gene regulatory network [20]. Note that for different species or measurement types, the respective parts of the simulation procedure below can be adapted to account for prior knowledge on species specific network structures and alternative measurement models.

First, we defined synthetic gene regulatory networks resembling the structure of those known for *E. coli*. We extracted networks comprising 20 genes from the E.coli transcription factor - gene network available at Regulon-DB (version 9.4) [20] preserving properties of the network graph by using the modularity-driven algorithm available in GeneNetWeaver [54] (see methods 4.1.1). We extracted 1000, 100 and 200 networks for respectively training, validation and testing of the classifiers with the configuration of GeneNetWeaver shown in section S1.

**Figure 2.**
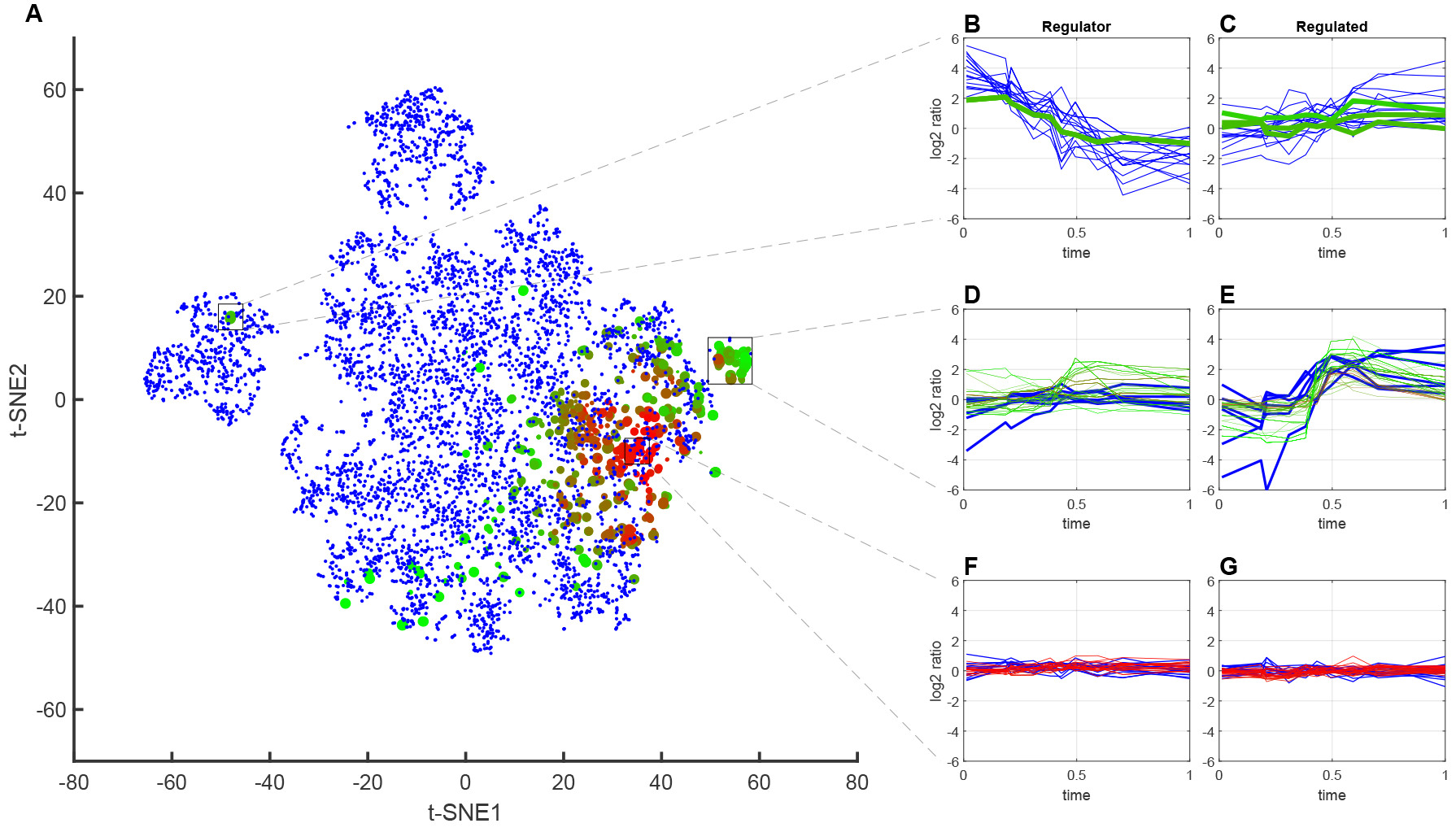
Synthetic data comprising dynamical behavior of regulated and regulating genes in experimental time series. Comparison of synthetic example dataset (*16* in Table S3) with measurement simulation (blue) and experimental data (green to red) of transcriptomic measurements of *E.coli* recovering from stationary phase (see more details in section 2.5). The color coding of the experimental data corresponds to the similarity of the time course to the synthetic regulator/regulated pairs from green (similar) to red (different). For details see *Maximum Mean Discrepancy witness function* in section 4.4.(A) t-SNE projection of concatenated time-courses of regulators and regulated genes. (B-G) time courses of transcripts inside the rectangular regions with the regulator in (B,D,F) and the regulated gene in (C,E,G) with the same color code as in the t-SNE projection. Similarity between synthetic and experimental data is high exclusively for gene pairs exhibiting an active regulation interaction.

Second, we generated synthetic microarray time-series data for these gene regulatory networks, under a variety of different perturbation conditions. To allow us to investigate what type of synthetic training data is best suited for reconstructing networks from a specific experimental dataset, we explored different types and extents of gene perturbation, initial conditions and measurement models. Specifically, we considered random dynamical models created individually for each network, thereby accounting for the uncertainty in their kinetic parameters (see section 4.1.2). We generated 30 datasets with different combinations of 1) numbers of genes affected per perturbation (single gene perturbed, multiple genes perturbed), 2) initial activation of perturbed genes (three normal distributions with *μ* = 0.5/0.4/0.4 and *σ* = 0.1/0.05/0.1) and 3) perturbation signal type (five settings of mixed, fixed, increasing, decreasing and pulse signals). Additionally, we had ten datasets with initial activation of perturbed genes sampled from lognormal distributions, and five for which all genes had the same perturbation signal applied. (See dataset configurations in Table S3.) For the single perturbation setup this resulted in mean 3.65 (standard deviation 2.41) perturbations per network. For the multiple perturbation setup we generated five perturbations per network with mean 3.89 (standard deviation 1.85) genes affected per perturbation. The resulting 264,503 dynamical systems were simulated until steady-state and we extracted ten time points distributed according to our experimental dataset (section 2.5).

We investigated the similarity of the simulated data and the later considered experimental *E. coli* time-series data by means of two-dimensional t-SNE projections [62] and quantitatively by Maximum Mean Discrepancy [57] (Fig. S1a). The features for both analyses were all pairs of regulator/regulated genes, concretely the concatenation of two time courses of length ten (resulting in vectors of length 20). For t-SNE, we selected 5000 random pairs from the synthetic dataset. From the experimental data we selected all regulators, but maximally five randomly selected regulated genes. The resulting projections show a varying degree of overlap, exemplified by dataset *16* (see Table S3) in Fig. 2a. There, the experimental data is color coded by the value of the witness function, yielding larger values where the distributions of synthetic and experimental data are more different (see 4.4 for details). The projections show clusters of distinct temporal behaviour of regulator and regulated transcripts (Fig. 2b/c), allow for the identification of single regions of low coverage of the experimental data by simulations (Fig. 2d/e) and highlight the presence of multiple experimental and synthetic regulator/regulated pairs with low log_2_ fold changes (Fig. 2f/g).

The overlap regions with high similarity comprise regulator/regulated gene pairs with higher *log*_2_ fold changes in contrast to those with lower similarity (Fig. 2f/g). This observation suggests that the higher similarity could be indicative for active genes. We investigated this relationship by comparing genes with high similarity to those reported to be active in the publication of the experimental dataset [53]. We evaluated the witness function for each experimental regulator/regulated pair (according to Regulon-DB) with an equal number of synthetic pairs and computed enrichments of the gene classes introduced in [53]. The comparison to the reported *activity scores* of the corresponding experimental conditions (*Early recovery in LB* and *Late recovery in LB*) for the example dataset yields a correlation of 0.45 (p-value 0.016, Fig. S1b) and indicates similarity between active experimental and synthetic pairs of regulators and regulated genes.

This relationship between regulatory active gene-gene interactions and similarity of experimental and synthetic time courses indicate that our simulations capture relevant experimentally observed dynamic behaviors.

### 2.2 Supervised learning of gene regulatory networks

The generated synthetic data comprises the time-series data, and the corresponding ground truth regulatory network. We utilized this for training supervised learning algorithms. We considered random forests and logistic regression, and explored hybrid convolutional recurrent neural networks; to date these have not been considered for gene regulatory network reconstruction. We used these algorithms as edge-classifiers, predicting the presence/absence of regulation between pairs or triplets of genes, represented by their transcripts’ time courses.

From the above synthetic data we filtered for only informative training data. Specifically, we created training sets by extracting groups of genes, i.e. 2/3-tuples of transcripts, that can be reached by the signal of a perturbation along the regulatory edges of the network (i.e. in the transitive closure of a perturbed regulator). We concatenated this set with an equally large set of transcript groups without any regulation (see methods section 4.2). The selected time courses were transformed to log_2_ scale and either used without further adaption (subsequently *DefRef* for *default reference*) or additionally augmented with a simple simulation of the experiment, emulating the log_2_ ratio to an unknown base level of expression (*SimRef* for *simulated reference*). In both cases we applied realistic microarray noise to the resulting data points (see methods section 4.1.5). We trained the classifiers and compared their test performance to state-of-the-art unsupervised methods, namely *GENIE3* [31], based on random forest regression to rank possible regulators for each individual gene, *dynGENIE3* [30], an extension of *GENIE3* for time-series, and *Context likelihood of relatedness* (CLR) [17], which predicts z-scores for the mutual information observed between genes considering a background distribution and Pearson correlation. See methods section 4.7 for a more detailed description.

To use neural networks as classifiers, we focused on two hybrid convolutional-recurrent architectures. The first architecture (*ccbld*) is a variation of the *convolutional long short-term memory deep neural network* (CLDNN) [52], which stacks two convolutional, two LSTM and one dense layer below a dense softmax output layer. The second architecture (*cr*) is a simplification of the first and combines one convolutional and one recurrent layer followed by the output layer. For the latter architecture we varied the size of convolutional and recurrent network layers, benchmarking in total five different neural network models for all datasets and 19 neural networks for a subset (datasets *1,16,21,33,35,* see table S3). A full description is available in methods section 4.3.

The input data for the supervised learning approaches above was the simulated transcript groups described in the previous section. The output was the class of the individual edges between the genes in the respective groups of genes. The classes of edges considered were *no regulation*, *activation* and *inhibition*. We randomly selected 2.0 × 10^6^ training samples from each data set (or the whole set if its size was below 2.0 × 10^6^) in order to mitigate the effect of different training set sizes caused by the training data extraction of different perturbation setups. The different supervised learning models were trained as described in the Methods section.

**Figure 3.**
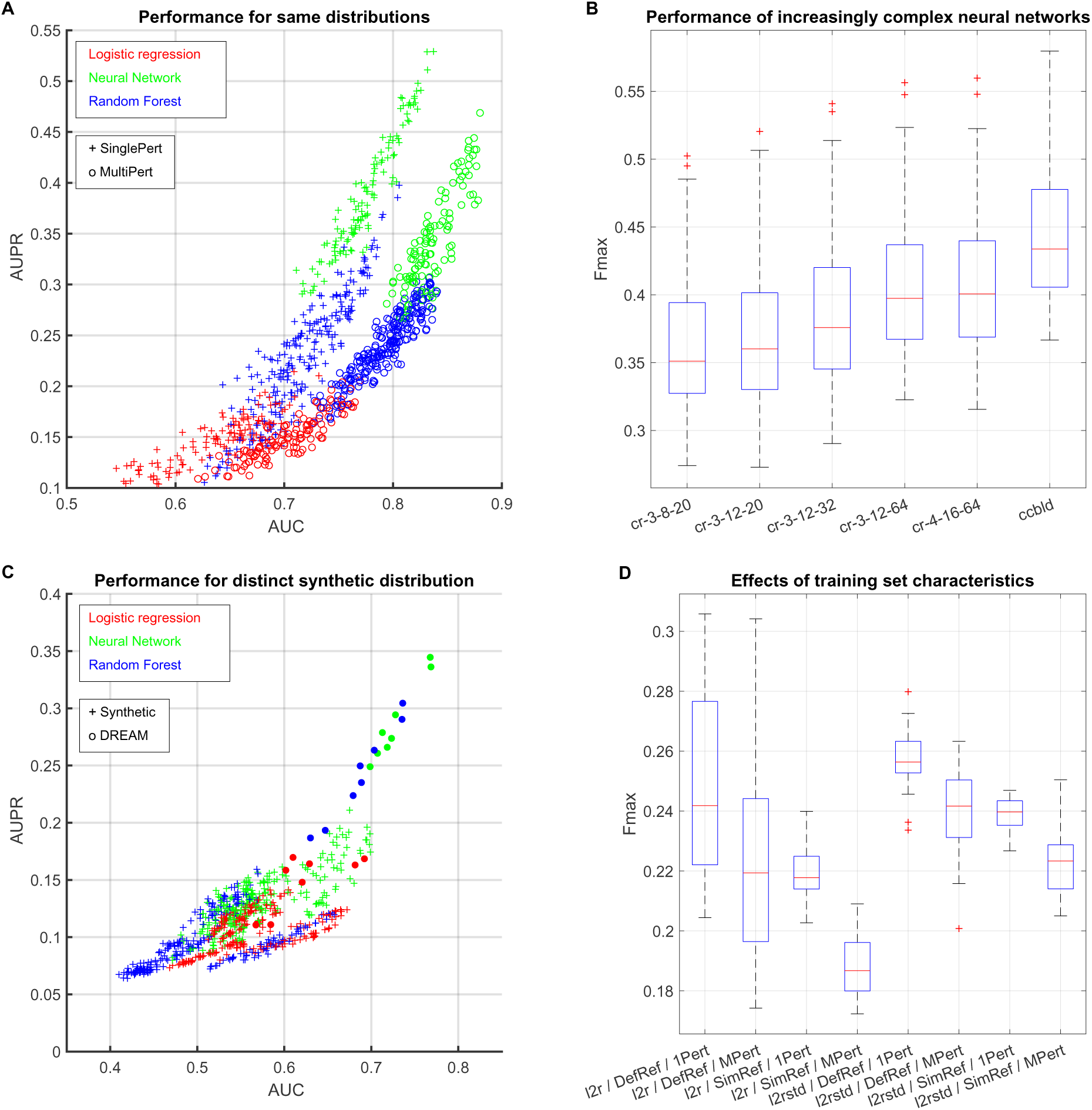
Network reconstruction results on synthetic datasets. **(A+B)** Reconstruction performance for test sets from the same distribution as the original training set of the classifier. **(A)** Area under ROC vs. area under Precision-Recall-Curve for the best classifiers of each type (neural network, random forest, logistic regression) for all synthetic testsets. The scatter symbol indicates whether the dataset had single (+, *SinglePert*) or multiple (o, *MultiPert*) perturbations applied. **(B)** Distribution of *F*_max_ scores for increasingly complex neural networks as described in section 4.3. (C+D) Reconstruction performance for DREAM4-like test sets. **(C)** Reconstruction performance for classifiers trained on 30 synthetic datasets (symbol +) compared to ones trained specifically on a DREAM4-like training set (symbol o). Area under ROC vs. area under Precision-Recall-Curve. **(D)** Distributions of *F*_max_ on validation set for different combinations of data pre-processing (*l2r* log_2_ ratio, *l2rstd* log_2_ ratio standardized per network), log_2_ ratio augmentation (*DefRef* default reference, *SimRef* augmented reference) and perturbation setup (*1Pert* single gene affected per perturbation, *MPert* multiple genes affected per perturbation).

### 2.3 Supervised learning on training and test data from the same distribution demonstrates superior performance of recurrent neural networks over simple classifiers

First we consider the ideal setting for the proposed supervised learning approach, where we precisely know the general mechanisms (e.g. kinetic model, interaction, and perturbation types) that give rise to the experimental data, but not the regulatory relationships between specific genes that we aim to reconstruct. y this situation by considering the same simulation settings for initial conditions, perturbation and measurement type for both training and test data sets, while idered regulatory networks differ between training and test data sets.

Throughout the manuscript we used the following metrics for reconstruction performance: *area under ROC* (AUC), *area under precision-recall-curve* (AUPR) and *F*_*max*_, the maximally achievable 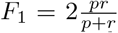 where *p* is the precision and *r* the recall at a certain threshold. Each of these values is computed per network in the test set and then averaged over the entire validation-or test set.

Overall, more complex models operating with larger feature vectors tend to achieve better results, we assume due to their capacity to learn the training distribution in more detail. The individual AUC and AUPR values per dataset for the most complex neural network architecture (*ccbld*), the most complex random forest (max. depth 100, max. features 150 %) and logistic regression with the best performing feature vector are shown in Fig. 3a. The two main sources of variation are the classifier type and the perturbation setup, defining groups of classifier/perturbation type pairs with differing performance. We observe that test data with multiple genes affected per perturbation yields lower AUPR, but higher AUC values (see also Fig. S2), which we assume to be due to the higher number of perturbations and higher coverage per perturbation as well as potentially several upstream regulator candidates per perturbation.

The best neural network classifiers perform consistently better than the best baseline supervised learning approaches. Additionally, we found that increasing model complexity in terms of neural network layers’ dimensionality improved the test reconstruction performance for five *cr* architectures with increasing number of internal nodes and the *ccbld* (Fig. 3b). These results indicate that more complex classification models may perform better where we have precise knowledge of the general mechanisms governing the experimental data.

### 2.4 Network reconstruction for distinct training and test data distributions demonstrates importance of realistic data simulation for learning

We next consider a more realistic setting for the proposed supervised learning approach, where we assume that we only approximately know the general mechanisms that give rise to the experimental data. y this situation by considering the different simulation settings for initial conditions, perturbation and measurement type for each training and test data sets, in ado having different regulatory networks between training and test data sets.

Specifically, as test data we followed the experimental time series setup of the DREAM4 challenge [43], with 200 networks, five perturbations per network and mean 6.64 (standard deviation 2.1) genes affected per perturbation. Each perturbation had a time-invariant activation, and we did not remove the perturbation signal after *t*_half_. We evaluated the reconstruction performance on this data for the classifiers trained on the original 30 synthetic training sets (non-DREAM4-like data) and compared it to results for classifiers specifically trained on a distinct training set of the DREAM4-like data. The classifiers trained on the DREAM4-like data outperform those trained on our original training sets (Fig. 3c), whose performance is decreased by 42.9 ± 14 % (mean ± std) *F*_max_ compared to the performance on their original training distribution (Fig. S3).

We assessed the network reconstruction performance for combinations of simulation/training settings, namely 1) application of data standardization, 2) augmentation of a log_2_ ratio reference and 3) single or multiple perturbations (Fig. 3d). The standardization of the inputs yields more consistent behavior across simulation settings compared to raw values. Within each group, using the default log_2_ ratio and single genes affected per perturbation was beneficial.

Despite generating the data with the same simulation model and the same parameters (e.g. measurement noise) or similar parameters (e.g. initial activation of perturbed gene) we observed a decrease in reconstruction performance compared to classifiers trained on the exactly same distribution. However, training data from the original 30 sets more representative of the target domain (e.g. without augmented log_2_ ratio references) or closer in its representation for the classifier training (e.g. data standardization) yield results comparable to logistic regression trained on the actual DREAM4-like data.

The results demonstrate the importance to our approach of training data appropriate for (or fine-tuned to) the subsequent test data for our approach.

### 2.5 Supervised learning achieves superior reconstruction performance for experimental time-series over state-of-the-art unsupervised learning approaches

We next evaluated different configurations of the simulations and classifiers in terms of their network reconstruction performance for experimental time-series data. Specifically, we analysed a time-series dataset measuring the transcriptomic responses of Escherichia coli (*E.coli*) recovering from stationary phase in rich media [53] available on Gene Expression Omnibus under accession GSE4363. For that, *E.coli* cultures had been collected and measured at eleven time points (0-1440 minutes) with cultures grown in Bonner-Vogel medium as a reference condition.

We constructed benchmark validation and test datasets from this data as follows. We used the first ten time points and extracted only time-series of transcripts with at most two missing values, each of which we interpolated linearly. The resulting set of genes was intersected with genes of the *E.coli* transcription factor gene network retrieved from Regulon-DB (version 9.4) [20], resulting in 1578 transcript species for analysis. We partitioned the remaining gene regulatory network of these transcripts, and split these partitions five times randomly into validation and test sets. For each of these ten sets we extracted 500 networks of size 20, which we performed the actual predictions on. The individual measurements of the original dataset were available as log_2_ ratios; alternatively we standardized these values for each sampled network of size 20 separately.

For each individual supervised classifier, we analysed which classifier configuration yielded the best results. Hyper-parameter evaluation resulted in best reconstruction performance for smaller models, both for neural networks and random forests. For neural networks, we compared *cr* architectures of different layer sizes and the *ccbld* architecture (Fig. 4a), and focused on the smaller three architectures (cr-1-3-8-20, cr-1-3-12-20 and cr-1-3-12-32) for further analysis. For random forests, the maximum depth of the trees and the type of the input feature vector showed an effect on the reconstruction performance (Fig. S4a), with tree depths between seven and thirteen and a feature vector of the concatenated raw time series *cas* as best configuration. For logistic regression, we used seven different input feature vectors (listed in Table 2) of which we identified the outer product of all absolute values (*oaa*) and the outer product of all absolute values combined with the outer product of all signed values (*oas.oaa*) as candidates for network reconstruction (Fig. S4b). We refer to this selected subset of neural network, random forest and logistic regression models subsequently as *selected classifiers.* Overall, we observe for the selected classifiers that random forests yield *F*_max_ values (0.279 +/− 0.041, mean +/− std) similar to neural networks (0.262 +/− 0.031) and better than logistic regression (0.192 +/− 0.015) on the validation set.

We studied the effect of different simulation and input configurations within the results of these selected classifiers. For neural networks and random forests separately, we assessed the effect of parametrization variants on the achieved *F*_max_ values with linear fixed effects models, which were selected according to the Bayesian Information Criterion (BIC) described in section 4.5. For neural networks (Table S5), we observe positive effects for the time-invariant perturbation signal (0.038, p-value = 9.8e-40), the augmentation of new log_2_ references (0.036, p-value = 1.2e-25) and standardizing the data (0.036, p-value = 1.6e-25), but not for applying the latter two jointly (-0.031, p-value = 4.06e-11). Moreover, perturbing multiple genes simultaneously had a negative effect when standardization was applied. For random forests (Table S6), the same overall effects are present, but with additional significant effects of the perturbation signals and gene initial activations. The results are reflected in the quartiles of the results grouped by the three main factors (Fig. 4b). For further analysis we only considered the identified beneficial parameter combinations (either augmentation with new references for the log_2_ ratios or standardization) and conclude that the time-variant perturbation signal settings are not necessary for our specific experimental dataset and subsequently only considered our sets *5,15,20,25,30,35* (see table S3), referenced as *final synthetic set*.

**Table 1.** Average reconstruction performance on the five experimental test set splits. Median and standard error bootstrapped from each of the five data sets individually and averaged subsequently. P-values determined from empirical H0 distribution of random structures. The concrete configuration of the supervised methods was chosen according to best performance on validation set.

| | AUROC | | AUPR | | $F_{\max}$ | |
| --- | --- | --- | --- | --- | --- | --- |
| | val ± stderr | avg. p-value | val ± stderr | avg. p-value | val ± stderr | avg. p-value |
| CLR | $0.52 \pm 0.005$ | 0.402 | $0.07 \pm 0.002$ | 0.429 | $0.15 \pm 0.002$ | 0.429 |
| dynGENIE3 | $0.51 \pm 0.003$ | 0.421 | $0.07 \pm 0.001$ | 0.422 | $0.15 \pm 0.002$ | 0.380 |
| GENIE3 | $0.53 \pm 0.003$ | 0.326 | $0.07 \pm 0.001$ | 0.431 | $0.16 \pm 0.002$ | 0.333 |
| Pearson | $0.48 \pm 0.004$ | 0.661 | $0.07 \pm 0.002$ | 0.484 | $0.15 \pm 0.002$ | 0.652 |
| Spearman rank | $0.48 \pm 0.004$ | 0.623 | $0.07 \pm 0.001$ | 0.431 | $0.15 \pm 0.002$ | 0.573 |
| Log. Regr. | $0.55 \pm 0.006$ | 0.166 | $0.10 \pm 0.003$ | 0.297 | $0.19 \pm 0.004$ | 0.156 |
| Neural Network | $0.58 \pm 0.009$ | 0.238 | $0.17 \pm 0.009$ | 0.370 | $0.27 \pm 0.009$ | 0.201 |
| **Random Forest** | **$0.63 \pm 0.009$** | **0.042** | **$0.20 \pm 0.010$** | **0.086** | **$0.34 \pm 0.010$** | **0.013** |

For classifiers trained on this final synthetic set, we verified consistent performance on experimental validation and test data. Average performance in terms of *F*_max_ over the five independent sets is highly correlated (Pearson correlation 0.967), but shows a bias toward better performance in the validation set (Fig. 4c). As a cause for this bias, we identified *experimental set 2,* whose validation set reconstruction worked much better than on the test set (Fig. S4c). In general, correlation between validation and test performance in the individual sets is lower and varies more (Pearson correlations 0.69, 0.43, 0.64, 0.65, 0.70) indicating a dependence on the partitioning of the original dataset. We also assessed the gain in reconstruction performance by utilizing the fully resolved time series information, i.e. by considering all ten time points versus only the first and the last one. We compared the existing random forests to ones specifically trained on the first and last time point (Fig S4d). For our *final synthetic set*, training/prediction on ten time points yielded on average 48.7 ± 16.3 % (mean ± std) better results than on two time points. For the second-best combination of simulation configurations (standardized data and using the default log_2_ reference) the advantage is 9.1 ± 6.2 % (mean ± std). The results indicate that the choice of a suitable simulation model allows for taking advantage of time-series information to significantly improve reconstruction performance.

Finally, we compared the reconstruction performance between supervised and unsupervised approaches. We selected the neural network, random forest and logistic regression configurations performing best on average over the validation sets of the five experimental splits, and compared each individual model’s performance on the test set to multiple state-of-the-art, unsupervised gene regulatory network inference methods. For our experimental dataset the supervised methods outperformed all unsupervised approaches in terms of AUC, AUPR and *F*_max_ (Fig. 4d, Table 1). For the supervised methods, random forest achieved the best results with an AUC of 0.65 and an AUPR of 0.22. These values are average results over 500 networks in each of five test sets. The *F*_max_ for individual networks vary between 0.012/0.007/0.008 and 0.53/0.97/0.97 for logistic regression, random forest and neural network (Fig. S5b).

In summary, our results demonstrate the feasibility and competitiveness of gene regulatory network reconstruction by supervised learning trained on synthetic data for transcriptomic time-series dataset, and identify beneficial configurations for simulation, data transformation and classifier training.

**Figure 4.**
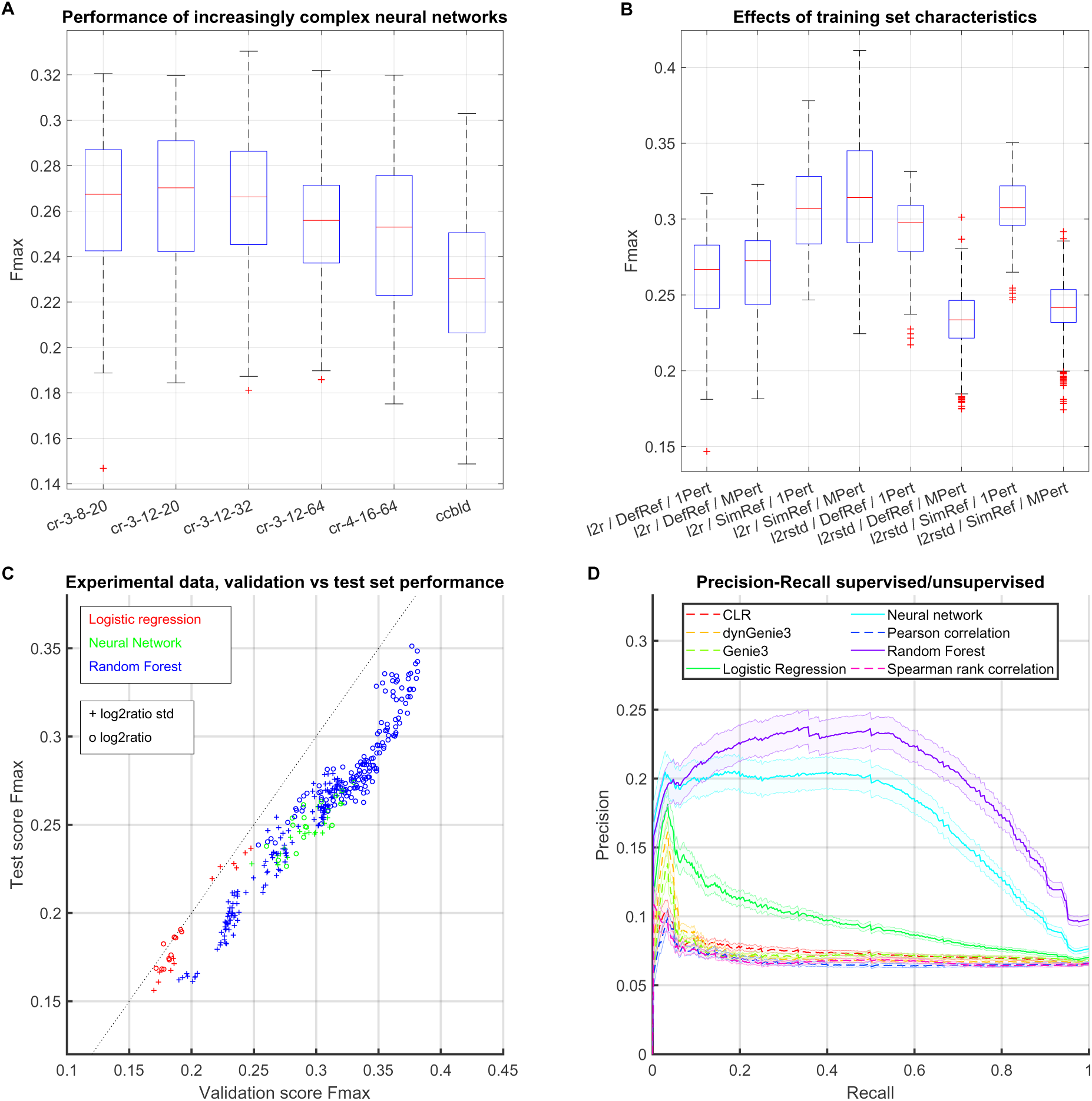
Network reconstruction results on experimental *E. coli* nutrient switch time-course data. **(A)** Distributions of *F*_max_ scores on validation sets for increasingly complex neural network architectures, as described in section 4.3. **(B)** Distributions of Fmax scores on validation sets for different combinations of data standardization (*l2r* log_2_ ratio, *l2rstd* log_2_ ratio standardized per network), log_2_ ratio augmentation (*DefRef* default reference, *SimRef* augmented reference) and perturbation setup (*1Pert* single gene affected per perturbation, *MPert* multiple genes affected per perturbation) for the *selected classifiers* (section 2.5). **(C)** *F*_max_ score for predictions of each considered combination of simulation-and classifier type on validation and test set. The scatter symbol indicates whether the dataset had single (+, *SinglePert*) or multiple (o, *MultiPert*) perturbations applied. **(D)** Precision-Recall curves for network reconstructions of the test set. The solid line is the mean over five test sets’ mean, each of which contained 500 networks. The shaded area represents the mean of the stderrs within each test set. For supervised methods, the selected parametrization was the one with the highest *F*_max_ score on the validation set.

## 3 Discussion

In this work, we present a surrogate learning approach for gene regulatory network inference from time-series data. We train classifiers on groups of genes and their time-resolved expression to detect patterns of regulation. The data for this training is exclusively synthetic and generated with a simulation model for transcription/translation, and a model for the measurement process. This approach allows for the generation of arbitrary amounts of training data and circumvents the need for experimental training data with correct labelling. The surrogate learning approach differs from conventional unsupervised reconstruction approaches by its use of mathematical modelling of biological and measurement processes; unsupervised approaches solve the inverse problem of inferring a model from a set of experimental observations. We consider models to solve the forward problem of simulating large amounts of representative synthetic data, train classifiers for structure learning on this data and subsequently predict directly on the experimental data.

We show good reconstruction performance for training and prediction on synthetic data from the same distribution and, a reduction of performance for closely related simulation or experimental settings. We found that less complex models coped best with the mismatch between the training and test data, presumably by introducing regularization, which lowers reconstruction performance on the training data, but yields better results for the relevant target domain. In contrast, we assume larger models learn to reconstruct gene-gene relationships from patterns specific to the training data distribution, transferring poorly to the target domain due to the lack of fit between synthetic training and experimental (or distinct synthetic) test data.

Transfer learning methods can mitigate a lack of fit between training and test data, and have been successfully applied to this end in other domains [16]. Indeed, we evaluated transfer learning for the neural network classifiers in two ways. First, initially training on exclusively synthetic data then subsequently re-training only the topmost neural network layers with experimental data and, second, by joint training with both synthetic and experimental data, differently weighted. However, neither strategy led to increased reconstruction performance on the experimental test set. We assume this is due to the small amount of experimental data, in particular after splitting in distinct training, validation and test sets.

Improvement of synthetic data generation can directly counteract the mismatch between synthetic and experimental data, and thereby beneficially impact network reconstruction. The data mismatch stems from modeling assumptions and formalizations [66]. While we systematically evaluated gene expression model parametrizations, perturbation variants and measurement models and standardizations, it is certainly possible to seek improvements by explicitly enumerating more simulation variants. It will be interesting to automate this process by considering generative models [24, 15] to learn *biologically representative* and *relevant* gene expression time-course patterns directly from experimental data. Considering the scarcity of experimental time-series data, training of such generative models could be augmented by synthetic data generated as presented in this study.

The presented *surrogate learning* approach required training of a large number of classifier instances. This bottleneck could be circumvented by defining suitable diversity measures of the simulated data, as well as similarity measures with the experimental data that are indicative for later reconstruction performance. To this end, we evaluated Maximum Mean Discrepancy, as a measure of similarity between data sets, and median pairwise distance, as indicator for the diversity within one dataset. Indeed, we see a trend of positive correlations between diversity and reconstruction performance and negative correlations between distance and reconstruction performance. However, those trends are masked by effects of distinct parametrizations of our data generation and show differences between the applied supervised classifiers (Fig. S6a). For further analyses explicit comparison of the relative similarity of two synthetic datasets to the experimental data could be beneficial [8].

Network reconstruction performance depends on the difficult to attain ground truth annotation of regulatory relationships. For the network reconstruction from the experimental *E.coli* data, we intersected all measured transcripts with those present in the current version of the gene regulatory network in the Regulon-DB and assumed the resulting network to be the ground truth for the evaluation of the reconstruction performance. It is conceivable that this ground truth set contains regulatory edges whose upstream genes were not active in the experiment, thus cannot be observed and lead to *false negative* predictions. The exclusion of such non-changing regulators, according to a differential expression analysis across time, could mitigate this issue and yield more accurate performance estimates.

The proposed reconstruction approach computes scores allowing for a ranking of potential gene-gene-interactions in an analyzed network, or for single genes of interest. Typically, thresholds for such rankings that achieve a desirable tradeoff between true/false positive/negative discoveries are derived from the optimal thresholds of a validation set. For the neural network classifiers, we applied the mean of the thresholds achieving the *F*_max_ in the validation set to the test set, and observed high correlation (0.87) and a decrease of 48.6 ± 8.7 % (mean/std) between *F*_max_ and the heuristically determined threshold (Fig. S6b). We expect improvements of such threshold estimate procedures through more precise modelling and taking into account prediction uncertainty. For instance, it will be promising to take advantage of the considered neural networks output that models the probability distribution over the three different regulation edge classes in the training data, allowing for informed choices of thresholds to evaluate the test data.

While we have focused here on transcriptomic time-courses and gene regulatory network inference, our study describes a generally applicable procedure to reconstruct different biological processes, e.g. signaling cascades, on the basis of different measurement techniques covering various biomolecules (e.g. RNA sequencing, mass cytometry) including single-cell measurements. Key for these applications is the availability of an appropriate simulation model of ‘generic kinetics’, concretely replacing the simulations from GeneNetWeaver [54], and a model of the measurement. While noise models for different measurement techniques are available [13, 1], we could not identify suitable biochemical models of ‘generic kinetics’ for biological processes other than gene expression [3, 67]. However, appropriate model classes [66] for many biological processes and concrete parametrized instances thereof [33] exist and could serve as starting point for generation of biologically representative data. While the classifiers and their configuration might be applicable to other bulk measurement data without further adaptations, single-cell data will entail an extension of the classifiers in order to operate on measurement distributions, instead of their bulk means. In summary, we expect surrogate learning to contribute a promising alternative to conventional network reconstruction approaches in a variety of systems biology applications in health and disease.

## 4 Materials and Methods

### 4.1 Simulation of representative training data

Our goal is the generation of biologically meaningful, synthetic data which is representative of microarray measurements. We divide this task in three steps: (1) The generation of genetic networks, which are small, but large enough to allow for non-trivial dynamics. (2) The simulation of intra-cellular transcription and translation based on generic biochemical kinetics. (3) The emulation of a microarray measurement process including noise and experimental setup.

We use and extend the software GeneNetWeaver version 3.1 [54] for network generation and simulation.

#### 4.1.1 Sampling of subnetworks

While our method aims to reconstruct entire gene networks, the working units of the algorithm are 2/3-tuples of genes. For diverse, generic behaviour, we extract these 2/3-tuples from simulations of networks of size 20, whose structure we extract from an actual biological network, specifically *E.coli’s* transcription factor gene interactions provided by Regulon-DB (version 9.4) [20]. For the synthetic training, validation and test sets we extract networks from the entire Regulon-DB graph, potentially resulting in overlapping network structures. However, each individual network is subsequently assigned individually sampled biochemical parameters.

We use GeneNetWeaver’s built-in functions to extract these networks from the Regulon-DB graph. This algorithm [42] randomly selects a seed gene from the source network and extends the graph iteratively by adding the neighbour whose addition maximizes the modularity of the new network. The modularity is here defined as the number of actual edges in the subnetwork minus the expected number in a randomized network with the same degree sequence. The procedure has been shown to preserve graph properties, such as motif enrichment, in the sampled sub-networks [42].

#### 4.1.2 Intra-cellular, biochemical simulation

Depending on the biological and experimental context, different mathematical models are suitable for simulation of representative data [23]. For RNA abundances in bulk measurements of cell populations, we explicitly modelled cellular abundances of RNA and protein, and protein-dependent production of RNA, mimicing the regulation by transcription factors. Furthermore we assume that stochastic fluctuations of gene activation, mRNA and protein concentration (such as bursts) at the single-cell level cancel out over the entire population and that Chemical Langevin equations (CLE) and Reaction Rate equations (RRE) are suitable for numerical simulations.

The biochemical model implemented in GeneNetWeaver consists of the following differential equations [43]:

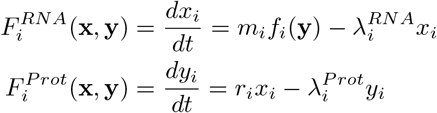

where 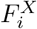 and 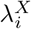 are the rate of change and degradation rate of component *X*, *m*_*i*_ is the maximum transcription rate, *r*_*i*_ the translation rate and x and y are vectors containing all mRNA and protein concentrations, with *f*_*i*_(·) denoting the relative activation of gene *i*.

The model of gene regulation is encoded in the activation function *f*_*i*_, which computes the mean activation of a gene *i* as a function of its transcription factors [43]. The underlying assumption is that the binding of the transcription factors is in quasi-steady state, which allows for the expression of the probability of combinations of transcription factors bound to the DNA and the explicit modelling of cooperative interactions including regulatory logic (AND, OR) [6, 43]. An example with two transcription factors is shown below:

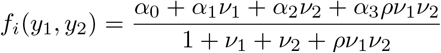

where *y*_1_, *y*_2_ are transcription factors, *α*_0_ is the basal activation of gene *i*, *α*_1_, *α*_2_, *α*_3_ are the activations for individual and both transcription factors bound and 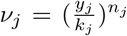 with dissociation constant *k*_*j*_ and Hill coefficient *n*_*j*_.

GeneNetWeaver uses a non-dimensionalized form of the system of the equations above [43], which bounds each state-variable between 0 and 1 and allows for easier, biologically meaningful random initialization of the biochemical parameters [67]. Additionally, transcription factors acting on one gene are randomly grouped in *regulatory modules*, whose members are randomly assigned to act as a complex or individually.

#### 4.1.3 Genes per perturbation and perturbation strength

Per perturbation we used two different ways to select the affected genes. In the setting *single* we generated one perturbation per gene, which had two or more downstream genes. The alternative *multi5* created a fixed number of five perturbations per network. Subsequently we determined the set of regulators *R*_1_ of genes with one downstream gene as well as the set of regulators *R*_2_ of genes with two or more downstream genes. For each perturbation and for each set, we sampled the number of genes *n*_*g*_ ~ *U*(0, |*R*.|) as well as (uniformly at random) which genes to perturb.

The actual perturbation strength *s*_*g*_ was computed according to

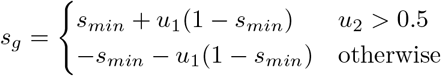

where *s*_*min*_ is the minimum perturbation of 0.5 and *u*_1_, *u*_2_ ~ *U*(0,1).

#### 4.1.4 Enhanced perturbation signal

GeneNetWeaver allows for single and multifactorial perturbations of different kinds (knock-down, knock-out, overexpression), each of which is implemented by a time-invariant change of the basal activation level *α*_0_ at *t*_0_ [54].

In order to represent more diverse dynamic behavior, such as cellular signaling, we replaced the previously constant perturbation signal by generic double-sigmoidal pulses, which allow for transient activation and deactivation. For this purpose we extended GeneNetWeaver with Gaussian-distributed basal activations for genes and pulse-like [10, 19] perturbation signals *s*_*g*_ (t) specific to each perturbed gene *g*:

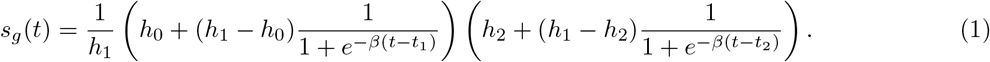

where *h*_0_, *h*_1_, *h*_2_ are the initial, intermediate and final amplitudes, *t*_1_, *t*_2_ are the half max times of the first and second sigmoidal transition and *β*_1_, *β*_2_ are the slopes of the transitions.

For the generation of datasets (see results) we parametrized *s*_*g*_ such that it creates pulses, increasing or decreasing sigmoidal curves or constants over time, where the latter reproduces the original behaviour of a fixed perturbation signal. Table S3 lists each dataset’s probabilities for choosing any of these four signal types and table S4 the configuration of the parameters of eqn 1 for each signal type.

#### 4.1.5 Measurement noise and experiment

The measurement noise was simulated with GeneNetWeaver’s built-in noise model for microarrays as originally developed in [61], which is implemented as multiplicative noise *x*_meas_ = *x*_sim_*e*^*w*^ and

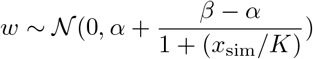

where *α* = 0:001, *β* = 0:69,*K* = 0.01 and *x*_sim_ is the simulated value.

The dimensionless output of GeneNetWeaver’s simulations represents the fraction of current RNA compared to the maximum steady-state abundance in linear scale. Our experimental dataset consists of log_2_ ratios between RNA measured under perturbation compared to a control. We mimic this behaviour by choosing a new reference point from the existing data points of a time course (assuming the transcript reaches a reference level during measurement), adding additional noise to the chosen reference and computing the *log*_2_ ratio. Concretely, we 1) sample the index of a new reference point in the synthetic data *i* ~ ***BB***(*n*, *α*, *β*) where *n* is the number of time points, *α* = *β* = 0.05 are the parameters of the beta-binomial distribution, 2) sample the new reference 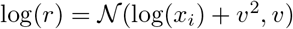 from a lognormal distribution with υ = 0:75 and 3) calculate the log_2_ ratio between the original simulation output and the new reference *r*.

### 4.2 Motifs and training data

Our neural network classifiers learn gene regulation patterns by analysing triplets of RNA abundances. Such network motifs, such as feed-forward loops, fulfill specific regulatory functions and have distinct enrichments in biological networks [4]. A known prior over this distribution of motifs could facilitate the inference of a genetic network. Random forests and logistic regression were performed on pairs of genes with input vectors created according to Table 2.

Training data is generated by perturbing one or several species in the networks of size 20 according to different perturbation patterns (see section 4.1.3). Since all species are initially in steady-state, gene-gene interaction is only apparent downstream of a perturbation. As training set *I*, we therefore only extract triplets *m* with at least one regulatory edge between the genes (set M_\0_) and with each species *s* either in the transitive closure *T* of the perturbation or having no edge *e* ∈ *E* at all.

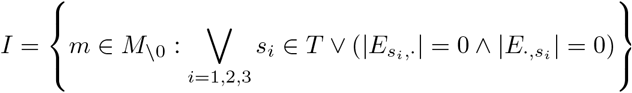

Subsequently we add sample pairs or triplets without any edge *M*_0_ independent of the transitive closure s.t. |M_0_| = |*I*| and add them to the training set.

**Table 2.** Names and descriptions of feature vectors used for random forest and logistic regression. All data was flattened to *n* × 1 vectors.

| Name | Description | Vector length (n) |
| --- | --- | --- |
| cas | Concatenated raw values | 20 |
| oas | Outer product of raw values | 100 |
| oaa | Outer product of absolute values | 100 |
| cas.oaa | Concatenated cas and oaa (see above) | 120 |
| cas.oas | Concatenated cas and oas (see above) | 120 |
| oas.oaa | Concatenated oas and oaa (see above) | 200 |
| cas.oas.oaa | Concatenated cas, oas and oaa (see above) | 220 |

### 4.3 Neural networks and motifs

For our study, we use neural networks operating on triplets of genes to learn patterns of gene regulation. We thus divide the problem of learning the edges of the entire directed graph *G* = (*N*; *E*) into the sub-problems of predicting individual edges in all 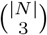 triplets of genes (network motifs [4]) in G. The predictions for each triplet are performed with the neural network based classifier resulting in |*N*| – 2 predictions per potential edge *E* = *N* × *N* in the final gene network. To combine these predictions, we take the mean over all motif-based edge predictions and furthermore - if applicable - the maximum over all perturbations for one individual edge.

We use hybrid convolutional and recurrent neural networks as classifiers and motivate this choice by the success of convolutions for feature extractions in other domains [39], as well as the explicit consideration of the sequential order of the time-series data by recurrent neural networks [28].

We consider (1) shallow architectures with one convolutional layer (convolution over 2-4 time points, dimensionality 8-32) and one recurrent layer (dimensionality 12-128), and (2) an adaption of the *convolutional long short-term memory deep neural network* (CLDNN) [52], which consists of two convolutional layers (3×3×256), one bidirectional LSTM (512), one LSTM (256) and one dense layer (512). In both cases there is a final softmax output layer for each individual edge in the motif. Subsequently, we refer to the latter architecture as *ccbld* or *ccbld-3-3-256-256-512-512* and to the smaller one as *cr* or *cr-p1-p2-p3-p4-p5-p6* with the following meaning of the placeholders *p*: (1) Convolution size over genes, (2) convolution size over time points, (3) number of convolutional filters, (4) number of dimensions in recurrent layer, (5) dropout fraction for recurrent layer and optionally (6) 0/1 indicating the presence of a direct link from the input data to the recurrent layer.

The input trajectories of the form x ∈ ℝ^3×*T*^, where *T* is the number of time points, are passed to the convolutional layer of size 3 × 3. For the large architecture (*ccbld*) the features extracted from the two convolutions are joined with the raw input trajectories and the resulting tensor is processed by a bidirectional LSTM layer. This returns an alternative representation of the data still containing the time series dimension. Both architectures output a fixed-length representation after the last recurrent layer, which is transformed using a fully connected layer. This is forked into the output layers with one hot encoded labels for each individual edge between the two involved genes A,B: 0 = no interaction, 1 = regulation of B by A, 2 = regulation of A by B.

We trained the neural networks with *RMSprop*, batch size 32, with 500000 random training samples per epoch, for a maximum of 100 epochs and early stopping (on validation loss) with a patience of 5 to 7 epochs and reduced the initial learning rates (*cr* 0.001, *ccbld* 0.0001) by 60 % after reaching a plateau (of validation loss) maximally twice.

The networks were built with the *keras* package (version 1.2.1) [12] using the *Theano* back-end (version 0.9.0) [59] and trained on NVIDIA Titan X and GeForce GTX 1080 Ti GPUs via the CUDA API [46].

### 4.4 Maximum Mean Discrepancy

In order to quantitatively assess the similarity between the synthetic and the experimental data sets’ regulator and regulated genes, we computed the Maximum Mean Discrepancy [26], concretely the estimator 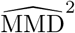 defined as

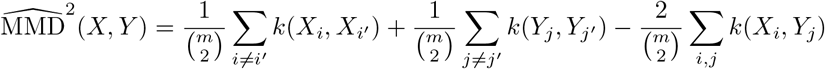

where *X* = {*x*_1_,…,*x*_*m*_},*Y* = {*y*_1_,…,*y*_*m*_} are samples from two different distributions (e.g. synthetic and experimental data) and *κ* is a kernel function, in our case the RBF kernel. The samples *x*_*i*_, *y*_*i*_ ∈ ℝ^20^ were the concatenated time courses of one regulator with one of its downstream genes. Per regulator we extracted maximally five regulated genes.

Following [57] we optimized the RBF kernel’s hyperparameter σ by maximizing the estimator of the t-statistic 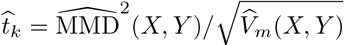 where 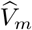 is the asymptotic variance of 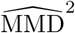(*X*, *Y*). We randomly partitioned the regulators in synthetic and experimental data into two sets, used one of them for optimization and the other for computation of the Maximum Mean Discrepancy with the optimized kernel bandwidth. We repeated this procedure ten times and report the mean values here. We performed these computations with a Python implementation^1^ provided for [57].

We assessed the similarity of individual regulator/regulated pair’s time courses by (the empirical estimate of) the *witness function* 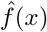 [26], whose magnitude indicates the difference between two distributions at *x*, evaluated with the optimized kernel bandwidth:

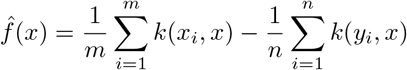

where *x*. were samples from the experimental data and *y*. were an equal number of randomly sampled regulator/regulated pairs from the synthetic data.

### 4.5 Linear fixed effects model for simulation/data parameters

To assess the effect of the parametrization of our synthetic training data, we sought to explain the *F*_max_ values, achieved by different classifiers on the experimental datasets, as linear model of the main factors: *A0Init* (initialization distribution of perturbed gene activation), *Sig* (type of perturbation signals used), *Stdize* (whether the input data was standardized per network), *AugRef* (application of randomly sampled reference for log_2_ ratio) and *MulPert* (multiple genes perturbed at once).

Starting from a maximal model containing all possible interaction terms (*F*_max_ ~ *Stdize* ∗ *AugRef* ∗ *MulPert* ∗ *Sig* ∗ *A0Init*), we used Matlab’s *stepwiselm* function for stepwise trimming of the terms according to BIC. The resulting significant coefficients (at *α* = 0.05) as well as BIC and *R*^2^ are shown in tables S5 and S6.

### 4.6 P-values and standard errors of AUROC/AUPR/*F*_max_

Following [56], we computed p-values for AUROC, AUPR and *F*_max_ for each individual network in all test sets by (1) computing the respective statistic for 10,000 random predictions, (2) fitting an exponential model to the obtained histogram and (3) computing the p-value as the integral under the exponential model between the achieved score and one. For AUROC and AUPR we used the model proposed in [56]

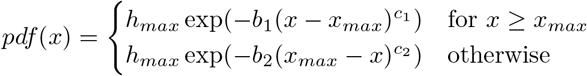

where *x* is the observed score and *h*_max_, *x*_max_, *b*_1_, *b*_2_, *c*_1_, *c*_2_ are the model’s parameters. For *F*_max_ we used the following model for the observed, exponentially decreasing histograms:

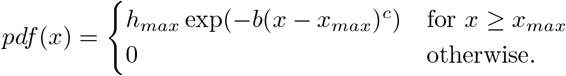

Standard errors of means for individual test sets of 500 networks each were estimated by bootstrapping (n=10000).

### 4.7 Comparison methods

The following methods each predict the edges of a gene regulatory network on the basis of statistical and/or dynamical models of gene expression directly from the experimental data, without prior training on training data.

#### 4.7.1 Context Likelihood of Relatedness

The Context Likelihood of Relatedness (CLR) [18] is a statistical approach based on mutual information between gene expression profiles. It extends the related relevance networks approach [9] by an *adaptive background correction* which computes a likelihood of an observed mutual information within its network context.

Under the assumption of a sparse interaction matrix, the distribution of all observed MI scores is assumed to be the background distribution, which is used to compute a z-score for a specific interaction’s MI score under an assumption of normality. This procedure is performed for both interaction partners and summarized as joint normal distribution 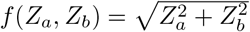 where *Z*_*a*_ and *Z*_*b*_ are the score z-scores for both involved genes. We ran the algorithm from the package CLR 1.2.2 in MATLAB 2017b with default parameters.

#### 4.7.2 GENIE3 and dynGENIE3

Gene Network Inference with Ensemble of trees 3 (GENIE3) [31] predicts regulatory interactions based on gene expression profiles by random forest regression on each gene independently.

The random forest approach covers interacting features and non-linear regulation and provides an importance measure for each regressor, specifically the total reduction of variance of the output variable induced by a split computed for tree construction. The importance measures from all regressions and genes pooled together are used as a global ranking for likely gene-gene interactions. We ran the algorithm from the package GENIE3 in MATLAB 2016 with default parameters.

Additionally, we applied *dynGENIE3* [30], an extension of *GENIE3* for time-series data, which explicitly models the temporal dependence of gene expression measurements with ODEs and finite difference approximation. We used the Matlab version^2^ on April 11th 2018 with default parameters.

## Footnotes

1 https://github.com/dougalsutherland/opt-mmd

2 Retrieved from http://www.montefiore.ulg.ac.be/huynh-thu/dynGENIE3.html

